# Characterization of chemotaxis in soybean symbiont *Bradyrhizobium diazoefficiens*

**DOI:** 10.1101/2025.10.14.682368

**Authors:** Matthew B. Lubin, Daniel H. Teixeira, Brittany J. Belin

## Abstract

Symbiotic relationships between nitrogen-fixing soil bacteria and legumes provide nearly half of all biologically fixed nitrogen on Earth, playing a crucial role in sustainable agriculture. These relationships rely on bacterial navigation of complex, dynamic soil environments to reach their plant hosts. Central to this behavior are bacterial motility and chemotaxis, the ability to sense and move toward host-derived signals in the rhizosphere. In the soybean symbiont *Bradyrhizobium diazoefficiens* USDA110, motility is controlled by dual flagellar systems, and this strain contains three putative but uncharacterized chemotaxis operons (*che1*, *che2*, and *che3*). Using targeted deletions of all three predicted *cheA* genes, we show that *cheA2* is the primary driver of chemotaxis toward soybean seed exudate in soft agar assays, and that the greater contribution of *cheA2* vs. *cheA1* in soft agar chemotaxis is due to its genomic context. Interestingly, we also found that *B. diazoefficiens* mutants that are incapable of chemotaxis in semisolid media retain wild type-like swimming speeds in aqueous media. These findings provide insight into how the agricultural inoculant *B. diazoefficiens* coordinates its chemosensory systems to respond to its host plant.

**IMPORTANCE:** Chemotaxis is crucial for the establishment of beneficial plant-microbe associations, yet mechanistic studies of chemotaxis have been limited to a handful of soil bacterial models, namely *Azospirillum brasilense*, *Sinorhizobium meliloti*, and *Rhizobium leguminosarum*. These three models represent only a fraction of the diversity found among plant- beneficial bacteria and agricultural inoculants. The soybean symbiont *Bradyrhizobium diazoefficiens* USDA110 is a commonly used soybean inoculant with exceptional nitrogen fixation efficiency, but the genetic control of chemotaxis in *B. diazoefficiens* has not been examined. Establishing *B. diazoefficiens* as a model of chemotaxis provides an opportunity to understand how multiple chemotaxis systems coordinate root colonization in this major agricultural symbiont and can enable comparative analyses of plant-microbe recognition strategies across agricultural bacteria.

## INTRODUCTION

Nitrogen-fixing bacteria are key players in the nitrogen cycle, converting atmospheric nitrogen into ammonia and thereby providing the building blocks for protein synthesis in all domains of life. In soil, symbiotic nitrogen-fixing strains known as rhizobia can provide ammonia directly to legume plant hosts (1). This symbiosis reduces the need for synthetic fertilizers and increases legume carbon sequestration, making rhizobia indispensable partners in sustainable agriculture. Given the high economic and environmental benefits of maintaining robust populations of rhizobia, it is important to understand how these microorganisms survive and navigate the soil habitat to establish beneficial associations with plants.

Natural soils are highly heterogenous and dynamic environments and relatively poor in nutrients, and soil-associated bacteria have thus evolved sophisticated motility and chemotaxis systems to enable them to sense and move towards nutrient gradients (5–7). Compared to bacteria in marine and animal environments, soil bacteria typically encode much higher copy numbers of chemotaxis receptors and signaling genes (8), suggesting that integrating multiple parallel chemosensory systems increases fitness in the soil niche (9–12). The relative contributions of individual chemotaxis gene networks have been examined in three model rhizobia, *Rhizobium leguminosarum*, *Sinorhizobium meliloti*, and *Azospirillum brasilense* (11, 13, 14). These studies revealed that the molecular functions of chemotaxis networks can be organism-specific, and they do not always contribute directly, or exclusively, to chemotaxis. For example, chemotaxis networks also can regulate diverse metabolic and cellular pathways (15) such as biofilm formation, exopolysaccharide production (16), nitrogen metabolism (17, 18), and cell division (19), as well as chemotaxis-independent aspects of colonization and nodulation (18, 20–25). Thus, it is difficult to predict whether individual chemotaxis gene orthologs actually regulate chemotaxis *a priori*.

Soybean is the most globally dominant legume by production volume, and it forms symbiosis with the rhizobium *Bradyrhizobium diazoefficiens* USDA110 (11, 26, 27). This strain possesses enhanced motility (2, 27–31) driven by its two distinct flagellar systems: a subpolar flagellum that drives swimming through water-filled pores, and multiple lateral flagella that facilitate movement across surfaces and through viscous environments (32, 33). Though regulation of flagellar genes in B. *diazoefficiens* has been shown to interact with metabolic and environmental sensing pathways (34, 35), and this strain can move directionally toward soybean exudate compounds (36–38), the genes required for *B. diazoefficiens* chemotaxis are unknown.

Canonical bacterial chemotaxis requires a signal transduction pathway that centers on a phosphorelay system involving methyl-accepting chemotaxis proteins (MCPs), the histidine kinase CheA, and the response regulator CheY (39–41). Analysis of the *B. diazoefficiens* USDA110 genome revealed 36 MCPs and three chemotaxis operons, each containing its own *cheA* gene (11, 42, 43). Here we use clean genetic deletions to analyze the role of each *cheA* genes in chemotactic signaling and motility control, as well as the relevance of their genomic positioning. This study establishes *B. diazoefficiens* as a new model for chemotaxis in soil bacteria, which may provide a foundation for engineering improved *Bradyrhizobium* spp. inoculants in agriculture (5, 44–46).

## METHODS

### Generation of deletion strains

We generated clean deletion mutants of *cheA1* (*bll0393*), *cheA2* (*blr2192*), and *cheA3* (*blr2343*) and combinations thereof using homologous recombination (47). For each targeted deletion, we designed DNA constructs containing 1 kb of sequence homologous to regions immediately upstream and downstream of the target gene. These sequences were synthesized by Twist Bioscience and cloned into the suicide vector pK18mobsacB, which carries kanamycin resistance and sucrose sensitivity markers. Full vector sequences can be found in **Fig. S4.**

The resulting plasmids were introduced into *Escherichia coli* S17 cells by electroporation using a Bio-Rad Gene Pulser (2.5 kV, 200 Ω, 25 μF). Transformed *E. coli* S17 cells were then used as donors for conjugative transfer of the deletion constructs into *B. diazoefficiens* USDA110. Conjugation mixtures were plated on selective AG medium plates (see (48) for recipe) containing carbenicillin (150 μg/mL) and kanamycin (200 μg/mL). Kanamycin-resistant colonies were then grown in non-selective media and plated on AG containing 5% sucrose for counterselection. Candidate deletion mutants were screened by PCR using primers flanking the deletion site and then confirmed by Sanger sequencing (Azenta Bioscience).

### Soybean seed exudate preparation

Soybean seeds (Deer Creek, Hutchison) were surface sterilized by rinsing three times with 90% ethanol, soaking for five minutes in a 5% bleach solution with mild agitation, and then rinsing five times with nanopure water. Exudate was prepared by soaking 100 g of surface-sterilized soybean seeds in 100 mL of double-distilled water for 24 hours at room temperature without agitation. The resulting aqueous solution was separated from the seeds and lyophilized at -80°C. The dried material was then reconstituted in 10 mL of nanopure water, yielding a 10X concentrated solution. This solution was sterilized by passage through a 0.2 μm membrane filter, flash frozen with liquid nitrogen, and stored in aliquots at -80°C for future use.

### Reverse transcription quantitative PCR (RT-qPCR)

Colonies on AG plates were used to inoculate 5 mL rich AG medium starter cultures and grown to exponential phase (OD_600_ = 0.45-0.7) at 30°C with shaking at 250 rpm and then were back-diluted to OD_600_ = 0.02 and grown again to exponential phase. Five mL cultures for each experimental condition were separated into 15 mL conical tubes and returned to the incubator for 8 hours with either 500 μL of 10X sSE (described above) or water controls. After incubation, total RNA was extracted using the RNeasy Mini Kit (Qiagen) according to manufacturer’s instructions, including initial lysozyme step and on-column DNase I digestion. First-strand cDNA synthesis was performed using 500 ng total RNA (quantified by NanoDrop, Termo Scientific) with AMV Reverse Transcriptase (Promega) and random primers in 20 μL reactions following the Promega recommended protocol. RT-qPCR was conducted using Power SYBR Green Master Mix (Applied Biosystems). Each 20 μL reaction contained 10 μL 2× SYBR Green Master Mix, 0.2 μL each of 10 μM forward and reverse primers, 1 μL diluted cDNA (1:20), and 8.6 μL nuclease-free water. Thermal cycling was set according to manufacturer’s instructions using a CFX Opus 96 Real-Time PCR System (Bio-Rad). Primer sequences can be found in **Table S1**.

### Growth curves

Colonies of *B. diazoefficiens* USDA110 strains on AG plates were used to inoculate 5 mL rich AG medium starter cultures and grown to exponential phase (OD_600_ = 0.45-0.7) at 30°C with shaking at 250 rpm. Prior to transferring to 96 well plates, cultures were then diluted to OD_600_ = 0.02 into rich AG medium (1 g/L arabinose, 1 g/L gluconate, 1/g Bacto yeast) or minimal AG medium (0.5 g/L arabinose, 0.5 g/L gluconate). For sSE treatment, media was supplemented with a 1:10 dilution of 10X soybean seed exudate (described above) or an equivalent volume of distilled water. 200 μL aliquots of each bacterial genotype were transferred to individual wells of a 96-well plate (Corning) in five wells per strain and condition. To prevent evaporation, wells were overlaid with 30 μL of mineral oil.

Growth was monitored using a Tecan Infinite M Plex microplate reader maintained at 30°C with continuous orbital shaking. OD_600_ measurements were recorded every 2 hours for 96 hours. Between readings, the plate was shaken at 180 rpm with an orbital amplitude of 2 mm to ensure adequate aeration. Growth curves were generated by plotting the mean OD_600_ values against time, with shading representing the standard deviation of three biological replicates.

### Semisolid agar chemotaxis assays

AG medium with very low carbon (0.1g/L arabinose, 0.1g/L gluconate) and 0.3% agar was autoclaved and 50 mL aliquots were poured into 12-cm square plates (Greiner Bio-One). Bacterial starter cultures were grown to early exponential phase (OD_600_ = 0.15-0.20) in minimal AG medium (0.1g/L arabinose, 0.1g/L gluconate), and 10 μL of culture was added to the center of each plate. Two sterile Whatman filter paper discs (6 mm diameter) were placed 4 cm away from the inoculation site on either side. One disc was saturated with 50 μL of concentrated soybean seed exudate (sSE), while the control disc on the opposite side received 50 μL of sterile water.

Plates were incubated at 30°C in a humidified chamber. Bacterial population expansion was documented by capturing images every 24 hours for 7 days using a Panasonic Lumix DSLR camera mounted on a fixed stand. Expansion of the bacterial cell population was measured using ImageJ, and bias of the expansion towards sSE was calculated as the expansion distance from the center inoculation site towards sSE as a fraction of the expansion distance in both directions:

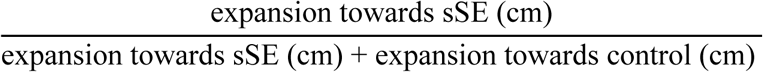

### Single cell tracking

Swimming of individual bacterial cells was analyzed using fluorescence microscopy. *B. diazoefficiens* cultures were grown to early exponential phase (OD_600_ = 0.2) in AG medium. The membrane-permeable nucleic acid dye SYTO9 (Thermo Fisher Scientific) was added to 1 mL culture to a 2.5 μM final concentration (1:2000 dilition from a 5 mM stock) and incubated with shaking for 10 minutes at 30°C. Immediately following staining, bacteria were spun down in a tabletop centrifuge at 750g for 8 minutes, gently resuspended in sterile saline (0.9% NaCl), and diluted to OD_600_ = 0.05. A 10 μL aliquot of the bacterial resuspension was placed onto a glass slide fitted with a SecureSeal™ Imaging Spacer and sealed with a coverslip.

Swimming cells were visualized using a Zeiss LSM 980 confocal microscope equipped with a 40X objective (NA 1.4) and kept in a temperature-controlled Pecon chamber at 30°C for the duration of the imaging. Images were acquired in fluorescence mode using 488 nm laser excitation and 500 nm emission with a frame rate of 17 frames per second. For each strain, multiple 30 second videos were recorded of different fields of view, capturing approximately 500 individual cell trajectories per condition. For chemotaxis experiments, cells were grown to early exponential phase and injected cells into a 𝜇 -Chemotaxis slide (ibidi) with either minimal AG media on both sides for homogenous control conditions, or with the right-chamber filled with media supplemented with 10X sSE, AG rich (1 g/L) media, or 10mM glutamate, following the protocol described in Ref (49). Briefly, the device is first overfilled with buffer free of chemoattractant or bacteria and the central channel’s ports were closed with plugs. Next, 65 µL was removed from one reservoir, replaced by 65 µL of chemoattractant-free bacterial solution, and then 65 µL of buffer was removed from second reservoir and replaced with bacterial suspension containing chemoattractant. All channel ports were plugged before microscopy observation.

Cell tracking and motion analysis were performed using a MATLAB script (Mathworks, MATLAB R2022b) adapted from Ref (50) to extract swimming trajectories and identify changes in swimming direction. Non-motile cells were removed from the dataset for analysis as determined by using a “stuck” validated by comparison to the calculated swim speeds of dead cells fixed in formaldehyde (Fig. S1, Table S1). Additional processing for the purposes of figure generation and subsequent statistical analysis of swimming parameters were written in Python. Statistical comparisons between strains were performed using Kruskal-Wallace tests, with significance defined as p < 0.05 (Python).

## RESULTS

### Organization and expression of chemotaxis genes in B. diazoefficiens USDA 110

Previous genomic analysis of *B. diazoefficiens* USDA 110 revealed three distinct chemotaxis operons: *che1, che2*, and *che3*. Each *che* operon contains an ortholog of the characteristic *cheA* histidine kinase and its response regulator *cheY*, suggesting each system could influence chemotaxis independently (**Fig. 1A**). Sequence analysis of each predicted CheA protein (CheA1, CheA2, and CheA3) indicates that CheA1 and CheA2 both possess the canonical domain architecture required for phosphorelay signaling and share 99% overall amino acid sequence similarity (**Fig. 1B**). CheA3 shares only 19% pairwise identity with CheA1 and CheA2 and lacks the characteristic CheY-binding domain required for downstream Che protein activation (**Fig. 1B**).

**Figure 1.**
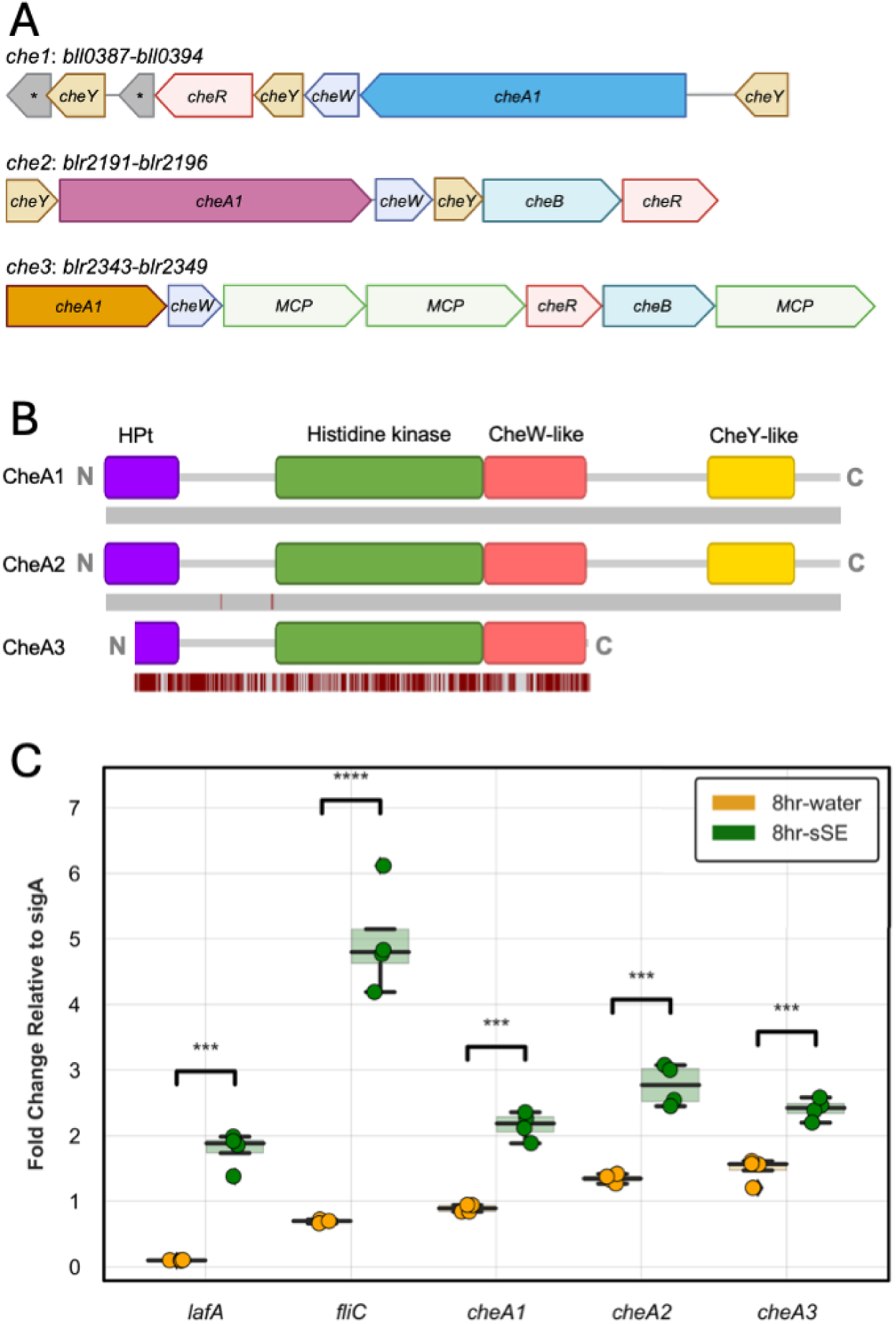
Chemotaxis genes of *B. diazoefficiens* USDA 110. **(A)** Gene architecture of three predicted chemotaxis operons, each with characteristic *cheA* genes named according to their operon number. Asterisks (*) connote uncharacterized ORFs. **(B)** Functional domain prediction of the encoded *cheA* proteins, overlaid with amino acid sequence misalignments (in dark red) compared to cheA1. **(C)** RNA transcript levels of genes coding for the primary flagellar component of the lateral flagella (*lafA*), the subpolar flagellum (*fliC*), and the three predicted *cheA* genes (*cheA1*, *cheA2*, *cheA3*) are upregulated compared to housekeeping gene *sigA* after 8 hours of incubation with soybean seed exudate (sSE) as compared to vehicle control (water). Significance determined by t-test: *** p < 0.0001, **** p < 0.00001

Prior studies in diverse organisms have shown that genes involved in chemosensation and motility are strongly induced by the presence of chemoattractants (16–18). We therefore tested whether each *cheA* gene is induced by seed exudate from soybeans (sSE), the native host of *B. diazoefficiens*. In liquid culture experiments, we found incubation of cells with sSE for 8 hours induced significant upregulation of the lateral and subpolar flagellin genes *lafA* and *fliC* as well as all three *cheA* genes (**Fig. 1C**).

### Swimming behavior of single chemotaxis mutants

To investigate the functional roles of each *cheA* gene, we generated scarless deletion mutants (Δ*cheA1*, Δ*cheA2*, Δ*cheA3*) via homologous recombination. We first assessed whether deletion of *cheA* genes affects general metabolic fitness. Growth curves comparing wild-type *B. diazoefficiens* with our panel of *cheA* mutants revealed nearly identical growth kinetics in AG rich media, with all strains exhibiting similar doubling times and reaching maximum optical densities after approximately 40 hours (**Fig. S1A**). Supplementation of cell cultures with soybean seed exudate (sSE) increased the maximum population density for all strains equally, with no differential effects observed among mutants (**Fig. S2A**). These results demonstrate that any subsequent differences in chemotactic behavior cannot be attributed to differences in growth dynamics in response to sSE.

We then examined colony expansion in semisolid (0.3%) agar plates containing minimal AG medium, a condition where expansion depends on swimming and on chemotactic navigation, as the cells at the inoculum site deplete minimal nutrients quickly during growth. Wild-type *B. diazoefficiens* formed characteristic circular expansion zones or “halos” of swimming cells that grew radially from the inoculation site, reaching approximately 5 cm in diameter after 7 days (**Fig. 2A**). Single deletion mutants Δ*cheA1* and Δ*cheA3* displayed expansion halos indistinguishable from wild-type halos, suggesting that neither CheA1 nor CheA3 is essential for motility under these conditions. In contrast, the Δ*cheA2* mutant exhibited significantly reduced expansion, reaching only 4 cm in diameter over the same period (**Fig. 2B**, p < 0.0001). This ∼20% reduction in expansion rate indicates that CheA2 plays a more critical role in coordinating swimming behavior compared to the other two CheA proteins on minimal medium plates.

**Figure 2.**
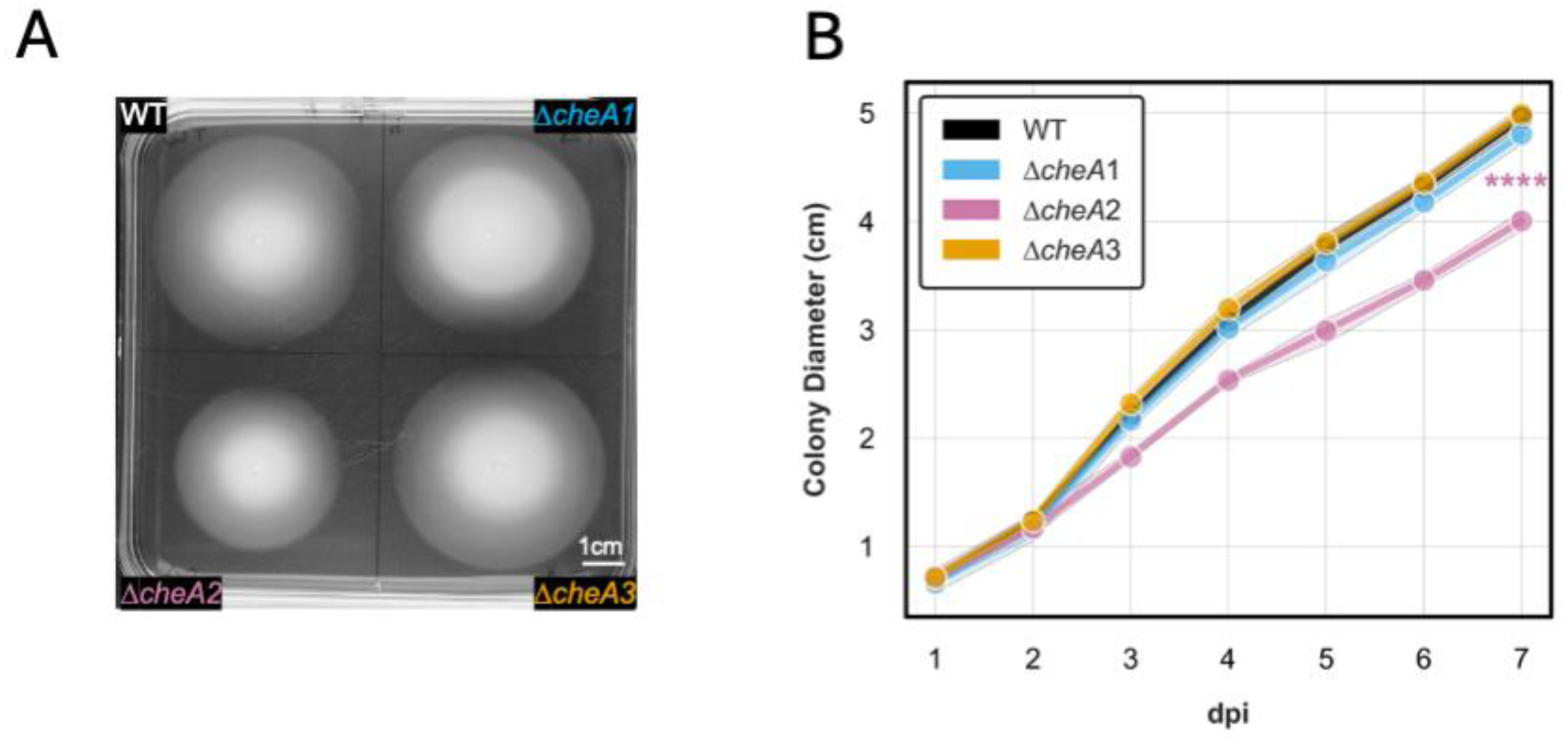
Loss of *cheA2* affects colony expansion in semisolid media. **(A)** Square semisolid (0.3%) agar plates were inoculated with cell suspensions of WT and *cheA* KO strains, which grew characteristic expansion halos, shown at 6 days post inoculation (dpi). **(B)** Daily measurements of colony diameter of each genotype show that Δ*cheA2* cells expand significantly less than WT or other single gene deletion strains (7dpi, p < 10e-6, ANOVA).

To highlight defects in chemotaxis specifically, we next examined bacterial expansion in a strong chemoattractant gradient. Filter discs saturated with concentrated sSE were placed on one side of the inoculation site, with water-saturated control discs on the opposite side, creating a directional attractant gradient across the plate. Wild-type *B. diazoefficiens* exhibited clear directional bias, with expansion halos extending preferentially toward the sSE source. By day 7, the halo of visible wild-type cells reached the sSE disc but not the control disc, with approximately 60% of their total expansion directed toward sSE (**Fig. 3A**). Both the Δ*cheA1* and Δ*cheA3* mutant strains displayed similar chemotactic bias to wild type (**Fig. 3A-D**), while Δ*cheA2* mutants showed a significant reduction in the directional bias of their expansion halos (**Fig. 3C**).

**Figure 3:**
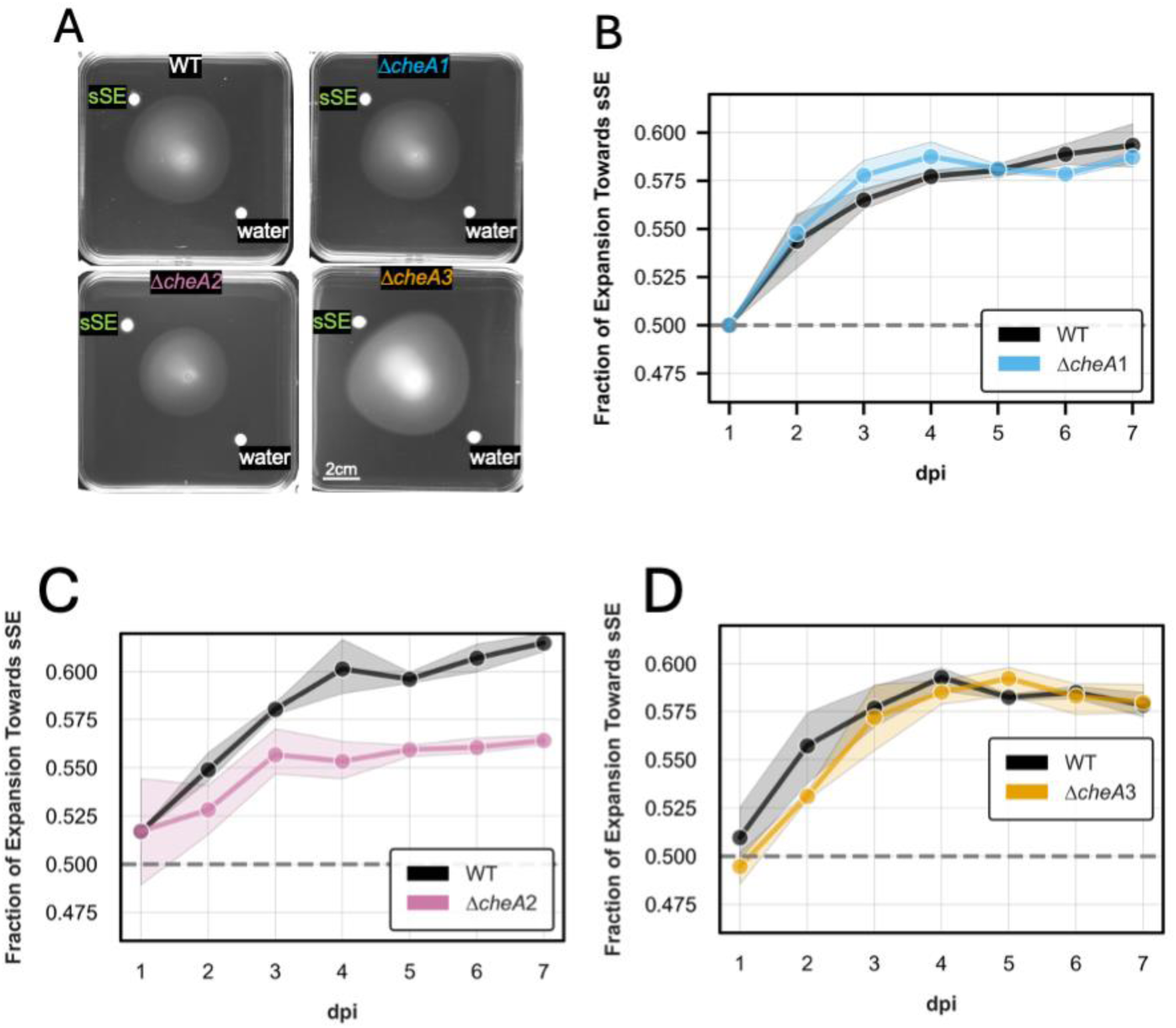
Loss of *cheA2* affects chemotaxis in semisolid media. **(A)** Cell suspensions were dropped onto the center of plates with equally distanced discs of either 10x soybean seed exudate (sSE) or water as control, shown here at 7 days post inoculation (dpi). **(B-D)** Daily measurements of the expansion bias shown in (A) with each genotype paired to WT cells inoculated on plates prepared the same day. Δ*cheA2* showed significantly less chemotaxis bias (p < 10e-4, ANOVA), while loss of *cheA1* or *cheA3* did not affect this phenotype (p > 0.05, ANOVA). Data in (B-D) represent the average and standard deviation from n=3 plates.

### Chemotaxis of double and triple *cheA* deletion strains in semisolid agar

Despite its reduction in colony expansion compared to wild type, the Δ*cheA2* mutant still retained measurable directional bias in expanding toward sSE, indicating that loss of *cheA2* alone does not completely abolish chemotaxis. This suggests functional redundancy with at least one of the other *cheA* genes. To investigate this potential redundancy, we generated double and triple deletion mutants and assessed their motility phenotypes. First, we repeated our growth curve experiments for the double and triple mutant strains to assess their general kinetics. As for the single mutants, we found that double and triple mutants had identical growth kinetics to wild type regardless of the presence of sSE in the medium (**Fig. S2B**).

In homogenous semisolid agar plates, the double mutant Δ*cheA1::*Δ*cheA2* exhibited a significantly more severe defect than the Δ*cheA2* single deletion, with expansion halos reaching only ∼2 cm in diameter after 7 days, representing a 60% reduction compared to wild type (**Fig. 4A-B**). Loss of *cheA3* yielded strain-dependent effects. The Δ*cheA1::*Δ*cheA3* double mutant resembled the wild-type strain (**Fig. 4C-D**), and the triple mutant Δ*cheA1::*Δ*cheA2::*Δ*cheA3* was likewise indistinguishable from Δ*cheA1::*Δ*cheA2* (**Fig. 4G-H**). However, the Δ*cheA2::*Δ*cheA3* double mutant expanded slightly more than the Δ*cheA2* single mutant (**Fig. 4E-F**, p=0.0107, hinting to a potential role of *cheA3* as a negative regulator of chemotaxis when *cheA1* is present.

**Figure 4.**
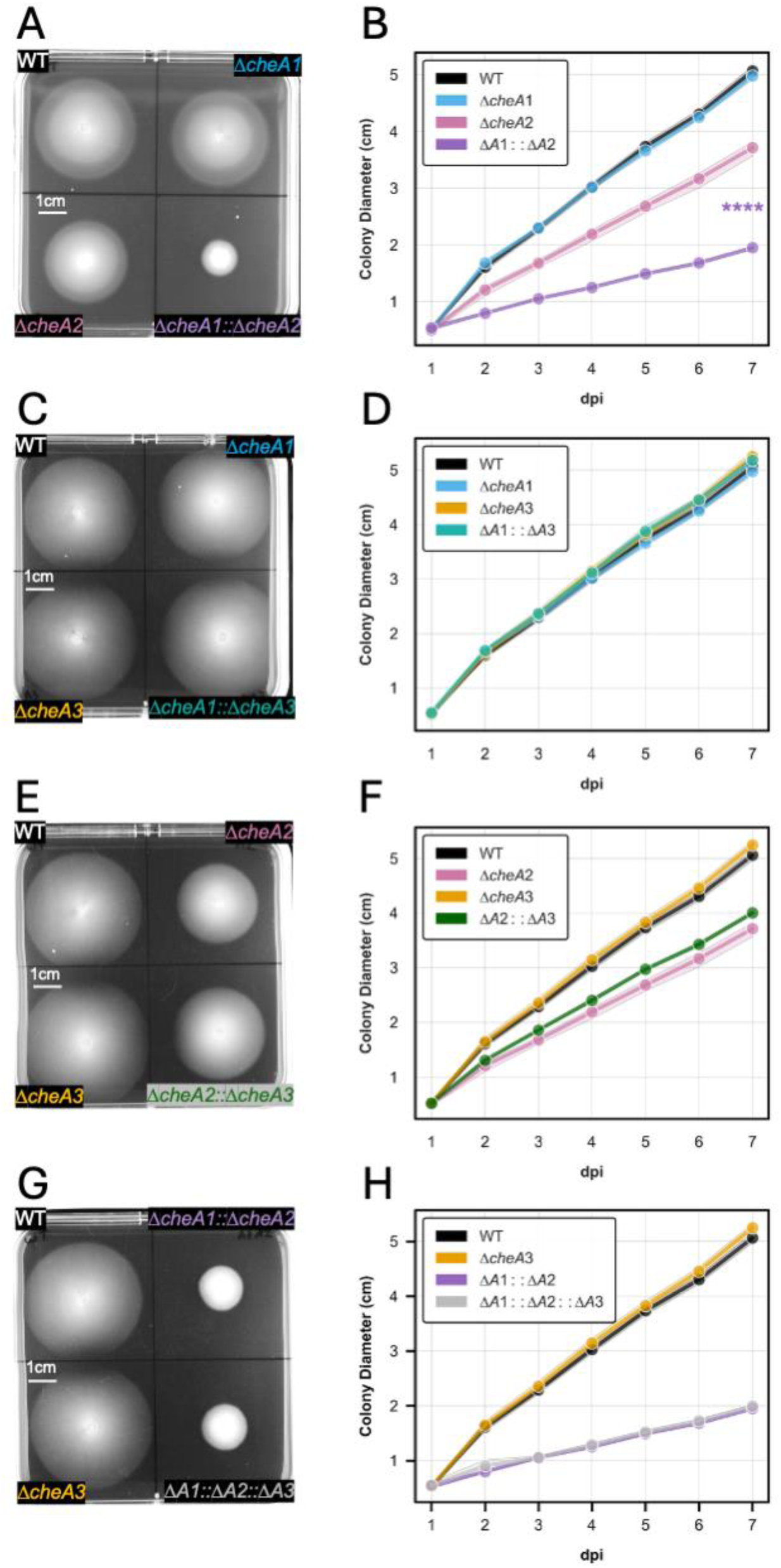
Double and triple *cheA* knockout motility. (A, C, **E)** Square semisolid (0.3%) agar plates were inoculated with cell suspensions of WT and *cheA* double-KO strains, along with two single gene KO strains, shown at 6 days post inoculation (dpi). **(B)** Daily measurements of colony expansions of each genotype show that Δ*cheA1::*Δ*cheA2* colonies expand significantly less than WT (p < 10e-12) and less than their Δ*cheA2* parent (p < 10e-6, ANOVA). **(D, F)** Daily measurements of cell expansion of each genotype show that the additional loss of *cheA3* does not reduce colony expansion, although Δ*cheA2::*Δ*cheA3* colonies expanded slightly more than Δ*cheA2* colonies (p=0.0107). **(G)** Inoculation of cells lacking all three *cheA* genes, shown at 6 dpi. **(H)** Daily measurements of colony expansion show that the additional loss of *cheA3* does not affect the Δ*cheA1::*Δ*cheA2* phenotype.

To investigate whether the motility defect of the Δ*cheA1::*Δ*cheA2* mutant reflected a loss of chemotaxis, we examined its behavior in the presence of a sSE gradient as we did for the single deletion strains. While wild-type *B. diazoefficiens* showed robust directional expansion toward the sSE disc, the Δ*cheA1::*Δ*cheA2* double mutant exhibited no directional preference in its expansion throughout the seven-day observation period (**Fig. 5**), indicating that this strain is effectively "chemotaxis blind" despite retaining residual motility. To confirm that the reduced expansion of Δ*cheA1::*Δ*cheA2* cells was due to loss of chemotaxis rather than loss of motility, we compared this strain to a previously described flagella-deficient mutant of *B. diazoefficiens* (32). The flagella-deficient strain showed no expansion from the initial inoculation point after 7 days in the presence of sSE gradients (**Fig. 5A**, bottom-right panel), confirming that expansion in these assays requires functional flagella. In contrast, the Δ*cheA1::*Δ*cheA2* double mutant maintained measurable expansion in both homogenous media and the sSE gradient condition, albeit without directional bias, indicating that this strain retains functional flagella and swimming capability. This effect indicates that *cheA1* and *cheA2* have an asymmetric redundancy: although *cheA2* fully compensates for a loss of *cheA1*, *cheA1* only partially compensates for the loss of *cheA2*.

**Figure 5:**
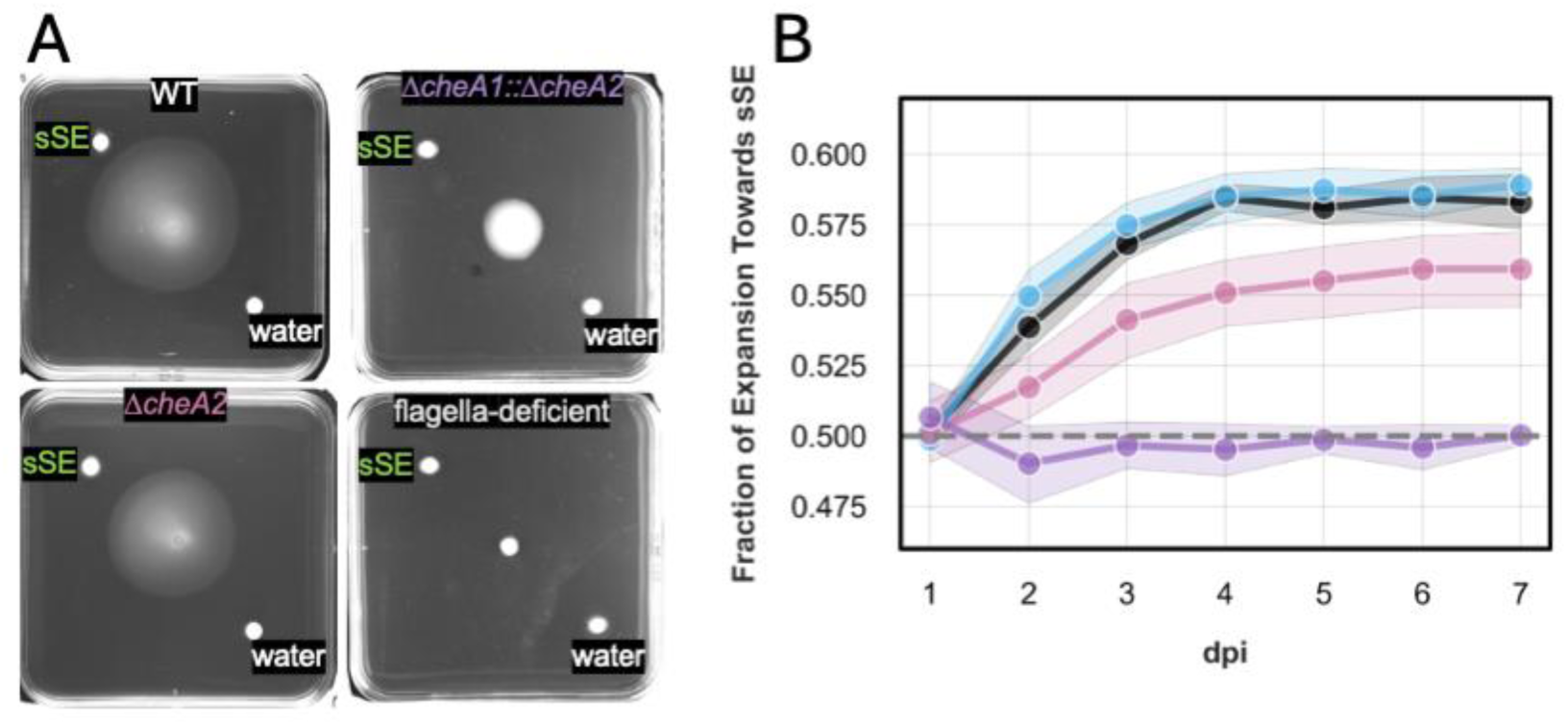
Δ*cheA1*::Δ*cheA2* shows no chemotaxis in semisolid media. **(A)** Semisolid (0.3%) agar plates with equally distanced discs of either 10x soybean seed exudate (sSE) or water controls were inoculated with WT, Δ*cheA2,* Δ*cheA1::*Δ*cheA2* cells, or cells from a strain lacking flagellar genes. **(B)** Daily measurements of the expansion bias shown in (A) paired with those of WT cells and Δ*cheA2* cells inoculated under the same conditions. Δ*cheA1*::Δ*cheA2* showed a complete loss of chemotaxis bias, significantly less than Δ*cheA2* single deletion (p < 10e-15, ANOVA). Data in (B) represent the average and standard deviation from n=3 plates.

Our finding that Δ*cheA1::*Δ*cheA2* cannot chemotax also suggests that *cheA3* does not promote chemotaxis. Plate chemotaxis assays on strains carrying *cheA3* deletions in the presence of sSE gradients support this interpretation, as deletion of *cheA3* did not affect expansional bias towards sSE, regardless of genetic background (**Fig. 6A-D**). The Δ*cheA1::*Δ*cheA3* double mutant displayed a directional bias toward sSE that is indistinguishable from both wild type and the Δ*cheA1* single mutant (**Fig. 6B**), while the triple mutant Δ*cheA1::*Δ*cheA2::*Δ*cheA3* remained completely chemotaxis-blind (**Fig. 6D**). Similarly, the Δ*cheA2::*Δ*cheA3* strain showed the same reduced chemotactic response as the Δ*cheA2* single mutant (**Fig. 6C**).

**Figure 6:**
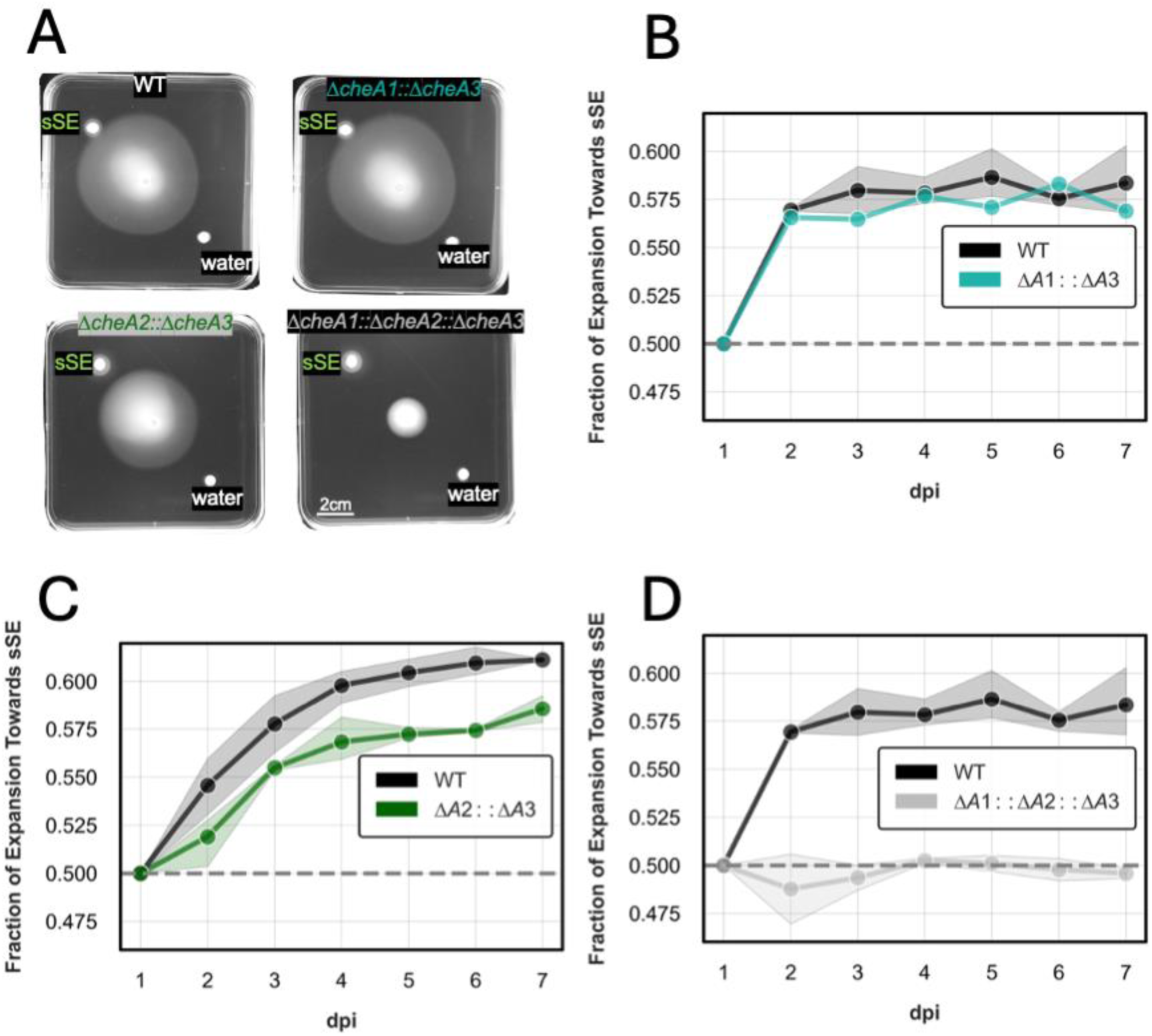
***cheA3* does not contribute to chemotaxis in semisolid media. (A)** Semisolid (0.3%) agar plates with equally distanced discs of either 10x soybean seed exudate (sSE) or water controls were inoculated with WT cells or cells missing *cheA3* in conjunction with the loss of one or both of *cheA1* and *cheA2***. (B, C, D)** Daily measurements of the expansion bias shown in (A) paired with those of WT cells inoculated under the same conditions. Data in (B, C, D) represent the average and standard deviation from n=3 plates.

### Motility of chemotaxis-deficient strains

To directly assess the motility of *cheA* mutant cells in aqueous environments, we performed single cell tracking analysis using fluorescence microscopy. All tested strains, including the Δ*cheA1::*Δ*cheA2* double mutant, displayed active swimming behavior and could reach swim speeds comparable to wild-type cells. Quantitative analysis of swimming speeds revealed from multiple experiments revealed no consistent differences between strains across biological replicates (**Table 1**). These single-cell observations confirm that the Δ*cheA1::*Δ*cheA2* strain retains fully functional flagella and normal swimming mechanics, supporting the conclusion that *cheA1* and *cheA2* govern chemotactic signal transduction and directional movement rather than flagellar motility *per se*.

**Table 1.** Loss of *cheA* genes does not reduce cell swimming speeds. . Average speeds of single cells of different *cheA* genotypes measured from six independent experiment (n>250 motile cell tracks per experiment). Superscript letters represent statistically indistinguishable groups (p > 0.01).

| **Exp #** | **WT** | ***ΔcheA1*** | ***ΔcheA2*** | ***ΔcheA3*** | ***ΔcheA1::ΔcheA2*** |
| --- | --- | --- | --- | --- | --- |
| **1** | 16.5 ± 4.9<sup>a</sup> | 17.1 ± 6.2<sup>a</sup> | 17.3 ± 5.1<sup>b</sup> | 16.0 ± 4.3<sup>a</sup> | 23.0 ± 8.0<sup>c</sup> |
| **2** | 19.8 ± 5.9<sup>a</sup> | 23.4 ± 8.7<sup>b</sup> | 17.1 ± 7.0<sup>c</sup> | 24.0 ± 9.0<sup>b</sup> | 20.5 ± 8.8<sup>a</sup> |
| **3** | 13.7 ± 4.0<sup>a</sup> | 13.7 ± 4.6<sup>a</sup> | 13.2 ± 4.6<sup>a</sup> | 13.7 ± 4.4<sup>a</sup> | 15.1 ± 5.3<sup>b</sup> |
| **4** | 14.1 ± 4.1<sup>a</sup> | 14.7 ± 4.2<sup>a</sup> | 13.6 ± 3.8<sup>a</sup> | 14.5 ± 3.5<sup>a</sup> | 15.8 ± 4.7<sup>b</sup> |
| **5** | 15.4 ± 4.2<sup>ab</sup> | 14.9 ± 3.9<sup>a</sup> | 15.9 ± 4.2<sup>b</sup> | 16.8 ± 4.1<sup>c</sup> | 29.6 ± 6.6<sup>d</sup> |
| **6** | 15.3 ± 5.5<sup>a</sup> | 16.0 ± 6.0<sup>b</sup> | 16.0 ± 5.5<sup>b</sup> | 15.9 ± 5.5<sup>b</sup> | 13.3 ± 4.5<sup>c</sup> |
| **AVG** | **15.8 ± 4.9** | **16.6 ± 6.0** | **15.5 ± 5.2** | **16.8 ± 5.7** | **19.6 ± 7.0** |

We also sought to determine the chemotaxis behavior of individual cells in aqueous media using established microscopy-based approaches. However, despite the robust chemotaxis we observed in soft agar assays, analysis of individual *B. diazoefficiens* cells in aqueous chemical gradients revealed no detectable chemotactic behavior. When exposed to gradients of sSE, rich AG medium, or 10 mM glutamate in microfluidic devices, wild-type cells displayed swimming patterns indistinguishable from control conditions (**Fig. S3**). Cell trajectories showed no directional bias, remaining radially symmetric regardless of attractant presence. Quantitative analysis confirmed that drift velocities along the gradient axis were centered at zero for all conditions, indicating no biased movement in either direction.

### Genomic context determines *cheA* functional hierarchy

Given that CheA1 and CheA2 share >99% amino acid sequence identity (**Fig. 1B**) yet display asymmetric functional importance, we hypothesized that their differential contributions to chemotaxis arise from their respective genomic contexts. To test this hypothesis, we generated a complementation strain in which the *cheA1* coding sequence was placed under the control of the *che2* operon promoter at the native *cheA2* locus (Δ*cheA2*::P*_che2_*-*cheA1*). Remarkably, this strain fully restored wild-type chemotactic behavior in semisolid sSE gradient assays, with expansion patterns and directional bias indistinguishable from the parental wild-type strain (**Fig. 7**). The complete functional rescue demonstrates that the *cheA1* gene product can fully substitute for CheA2 protein when expressed in the appropriate genomic context.

**Figure 7.**
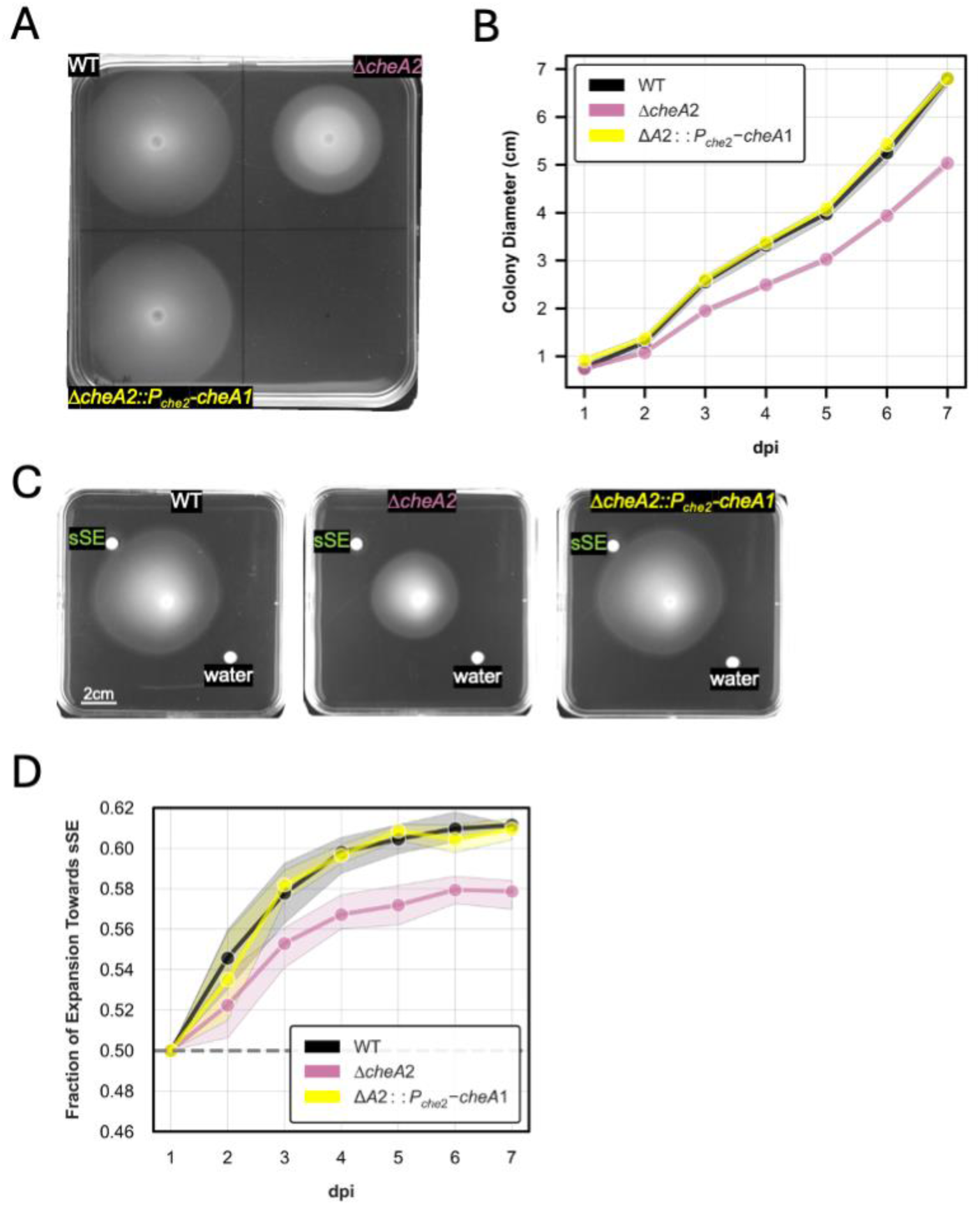
Complementation of the *cheA1* gene in the *cheA2* locus shows a reversion to WT- like phenotype. **(A)** Square semisolid (0.3%) agar plates were inoculated with cell suspensions of WT and Δ*cheA2* strains, along with a strain bearing the *cheA1* coding sequence at the *che2* operon locus generated in the Δ*cheA2* background (Δ*cheA2::P_che2_-cheA1*). Plates are shown at 6 days post inoculation (dpi). **(B)** Daily measurements of colony expansions shows full rescue of the expansion phenotype. **(C)** WT, Δ*cheA2,* and the complementation strains were used to inoculate plates containing equally distanced discs of 10x sSE or water controls. **(D)** Daily measurements of the expansion bias shown in (B) demonstrating the WT phenocopy of the complementation by *cheA1*.

## DISCUSSION

Robust chemotaxis is correlated with higher fitness in plant-beneficial soil bacteria and thus is an important consideration for engineering improved inoculants in agriculture. In this study we provide the first genetic analysis of chemotaxis in the soybean inoculant *B. diazoefficiens* USDA110. We find that, of the three *cheA* proteins encoded by this organism, only *cheA1* and *cheA2* contribute positively to chemotaxis toward soybean seed exudates in soft agar. While cells with deletions of *cheA1* or *cheA2* genes alone maintain some chemotactic ability, the combined loss of both genes abolishes directed movement toward sSE in soft agar assays. This suggests functional redundancy between the *che1* and *che2* systems in *B. diazoefficiens*, where either system alone is sufficient to mediate chemotactic responses toward host-derived signals.

Redundancy in the *che1* and *che2* systems may provide more robust chemotaxis in the native environment, ensuring that bacteria can still respond to host signals even if one pathway is compromised. However, our observation that expressing the *cheA1* coding sequence expressed from the che2 locus (Δ*cheA2*::P*_che2_-cheA1*) can restore wild-type chemotactic behavior highlights the importance of the operons’ differential gene regulation. We do not yet know what factors drive this differential regulation, nor the extent to which the relative induction of each operon is related to the nutrient sources available. Both AG medium and sSE both contain complex chemoattractant mixtures, and they do not capture the full milieu of nutrients encountered by native *B. diazoefficiens*, which includes compounds exuded at different stages of the soybean life cycle and from other plants, microbes, and insects in the environment. Future studies will be needed to determine whether differential *che* operon induction is tuned by specific compounds or whether it arises from their relative affinities for shared transcriptional regulators.

Though the absence of a chemotaxis phenotype for *cheA3* deletions in our assays could also stem from a mismatch between chemoattractants in AG and sSE and those in the native environment, we think this is unlikely. Unlike CheA1 and CheA2, CheA3 lacks the CheY-binding P2 domain essential for phosphotransfer to the response regulator. This structural difference precludes CheA3 from directly controlling flagellar rotation through the canonical phosphorelay pathway (51). Instead, CheA3 might function as a phosphate sink, sequestering methyl-accepting chemotaxis proteins (MCPs) and thereby attenuating signals from CheA1 and CheA2. Alternatively, CheA3 could be involved in adaptation mechanisms or serve regulatory functions unrelated to chemotaxis despite its genomic context. The slight but statistically significant (p=0.0107) enhancement of motility upon *cheA3* deletion in the Δ*cheA2* background supports a model where CheA3 competes with functional CheA proteins for MCP binding sites, though further biochemical studies are needed to test this hypothesis. Bacterial chemotaxis systems are known to include proteins that act to negatively regulate the cheA-mediated phosphorelay, and perhaps this truncated *cheA3* which is missing the CheY-like functional domain also serves as a chemotaxis response regulator.

A surprising finding in our study is that single *B. diazoefficiens* USDA110 cells do not chemotaxis effectively in liquid environments - at least not under the same conditions used to induce chemotaxis on soft agar or following protocols commonly used to induce aqueous single cell chemotaxis in *Vibrio* spp. and other rhizobia. This apparent discrepancy between our population-level chemotaxis assays and single-cell tracking experiments presents an intriguing puzzle. We propose three potential explanations. One possibility is that single-cell chemotaxis only occurs in response to specific, unknown chemotattractants not captured by our analysis or under conditions that are otherwise difficult to capture in the lab. Soil bacteria typically require more extensive starvation and a much narrower and lower concentration range to induce chemotaxis that biomedical model species. *B. diazoefficiens* may be on an extreme end of this continuum or only exhibit detectable single-cell chemotaxis on very long timescales.

Another possible explanation is that *B. diazoefficiens* USDA110 chemotaxis, at least when driven by *cheA2*, simply does not occur in liquid media. The physical constraints of moving through a semi-solid matrix might enable detection of chemical gradients through mechanisms that are inactive in purely aqueous conditions. There is already evidence that the *B. diazoefficiens* USDA110 flagellar motility systems are engaged in different environments, with the lateral flagella dominating in movement on or near surfaces. If the *che* systems of *B. diazoefficiens* USDA110 are primarily regulating lateral flagellar surface motility, this may not be evident in our aqueous chemotaxis system. Given that *B. diazoefficiens* USDA110 primarily resides adjacent to soil particles and plant tissues, it may be that robust chemotaxis through open aqueous environments makes an insignificant contribution to fitness in the native ecological context.

A final, theoretical possibility is that *B. diazoefficiens* exhibits emergent collective chemotaxis in liquid, where directional sensing emerges at the population level even when individual cells cannot effectively follow gradients.Formatting… Such phenomena have been observed in eukaryotic systems where cell clusters can achieve directional movement through mechanisms that do not require individual cells to sense chemical gradients, but a chemotaxis-like effect is nevertheless achieved over a long enough timescale (52–55). While the specific mechanisms would likely differ from eukaryotic collective chemotaxis, bacterial populations might achieve gradient sensing through cell-cell interactions or other collective behaviors that emerge only at high cell densities or at the longer timescales due to multiple generations spatially asymmetric cell divisions (56, 57).

Regardless of the underlying mechanism, our data indicate that chemoattractant responses of *B. diazoefficiens* USDA110 in liquid are distinct from previously studied soil bacterial models, providing the opportunity to examine the hypotheses. Future work examining the relative importance of these systems in more complex soil-like environments all will be important to understand how *B. diazoefficiens* chemotaxis systems are integrated in real-world agricultural settings. Understanding how *B. diazoefficiens* USDA110 responds to soils could benefit agriculture, including by optimizing the timing of inoculation relative to irrigation or development of improving carrier materials for bacterial inoculant delivery. Such improvements could enhance the robustness of plant-microbe interactions across varying soil conditions, an important consideration as climate change leads to more variable precipitation patterns in agricultural regions.

## AKNOWLEDGEMENTS

This work was supported by NIH grants R00GM126141 and R35GM147015 (to B.J.B) and Carnegie Institution for Science endowment funds. The authors thank A. R. Lodeiro (National University of La Plata, Argentina) for generously sharing flagellar-gene knockout strains of *B. diazoefficiens*, Birgit Scharf and Scharf lab members (Virginia Tech, USA) for training in rhizobium chemotaxis assays, and Mahmud Siddiqi (Carnegie) for microscopy assistance. We also thank members of the Belin lab for helpful discussions and feedback on the manuscript. We are grateful to Carnegie Embryology’s IT, front office, and facilities staff for making our work possible.

## SUPPLEMENTAL FIGURES

**Figure S1.**
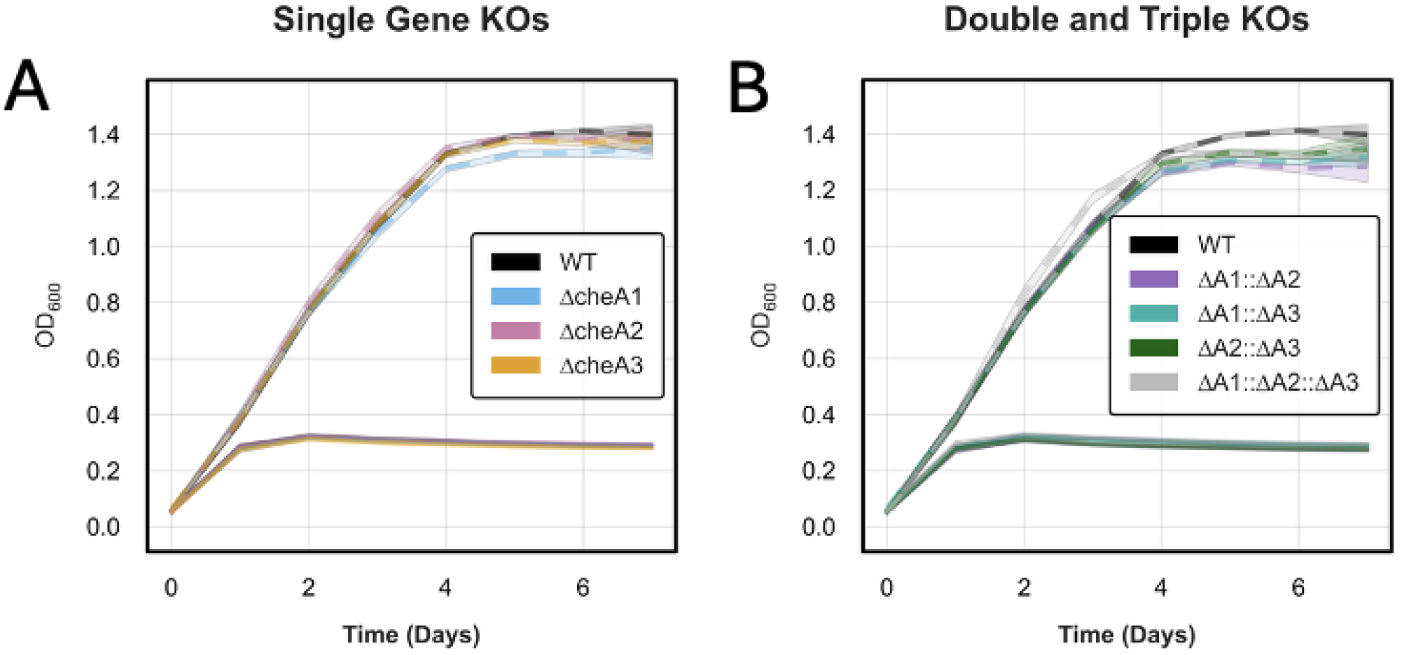
Loss of *cheA* genes does not affect growth. Liquid growth curve assays of WT cells compared to **(A)** single *cheA* gene knockout lines or **(B)** double *cheA* gene knockout lines grown in minimal AG media alone (solid line) or supplemented with soybean seed exudate (sSE, dashed lines). No significant difference was observed between the average growth rates or carrying capacity of any of the strains. Shading represents standard deviation from five technical replicates.

**Fig S2.**
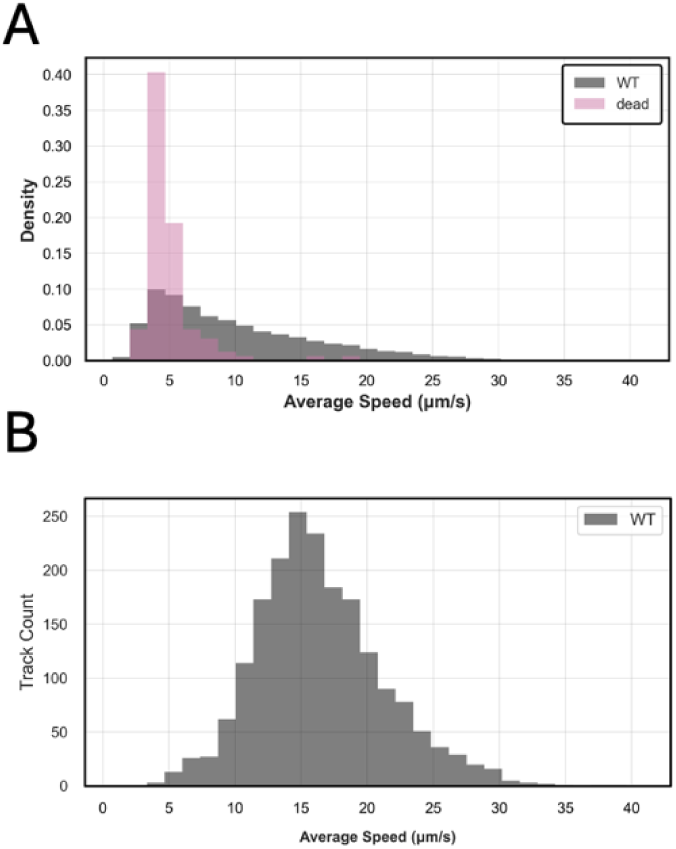
Non-motile cells can be mostly filtered out by using ‘stuck setting parameters.’ To account for only motile cells capable of swimming, a “stuck determination” was used such that if a cell moved less than 2 micrometers over the course of 6 consecutive frames at any point, it was filtered out. **(A)** Unfiltered dataset shows a high proportion of identified tracks of low speeds in both WT live cells and cells fixed in formaldehyde before staining. **(B)** After a ‘stuck’ cutoff parameter was applied, track speed distribution is normal. Filtering removed 92% of dead cells, but only 21% of live cells.

**Figure S3.**
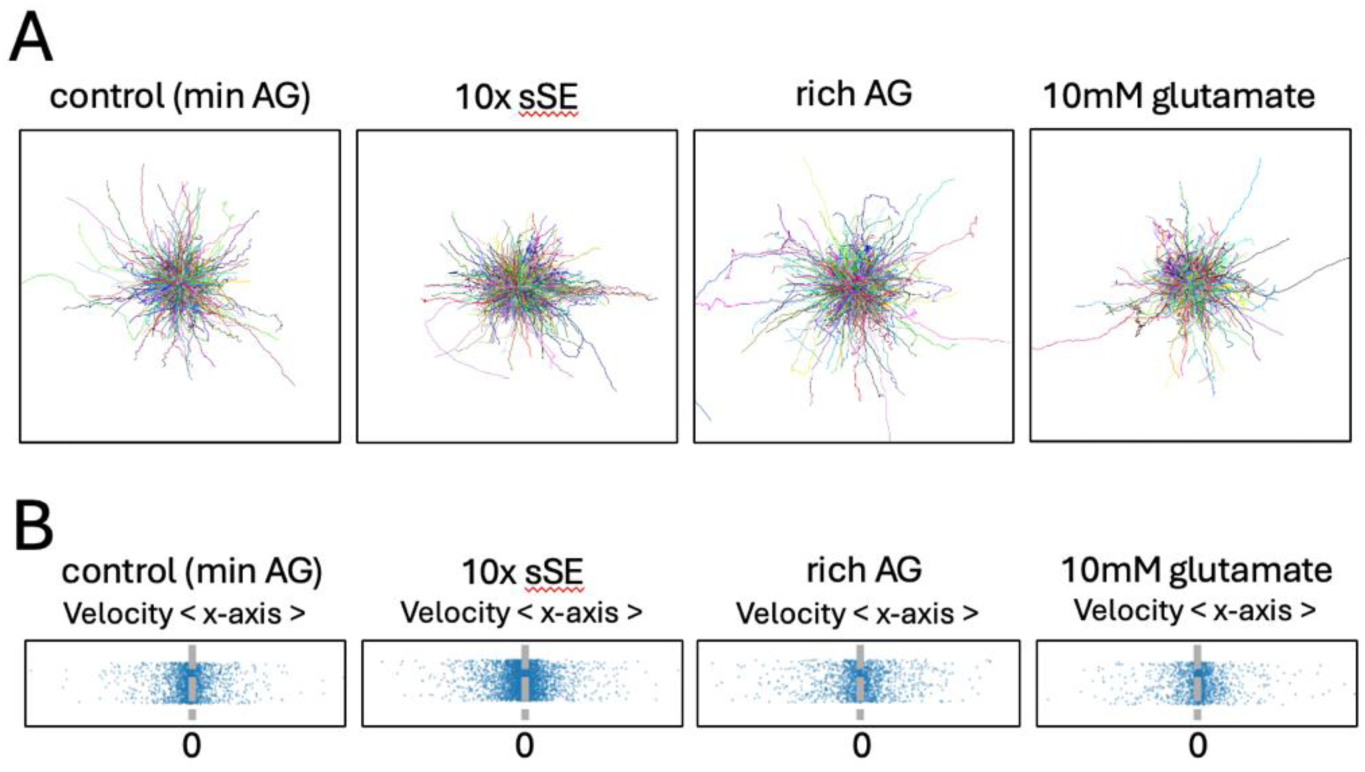
WT cells do not demonstrate a swimming bias across a gradient of potential chemoattractants. **(A)** RosePlots of 500 WT cell swimming trajectories from each condition, showing no directional bias in displacement of overall cell movements. **(B)** Drift velocity with respect to the x-axis, or ^𝑑𝑥^. Cells that swim equally fast in the positive (right) or negative (left) direction would have a net velocity of 0, which is the average drift velocity of cells in all conditions. For each condition, chemoattractant media was placed in the right chamber, such that the concentration gradient is highest on the right and lowest on the left.

**Figure S4.**
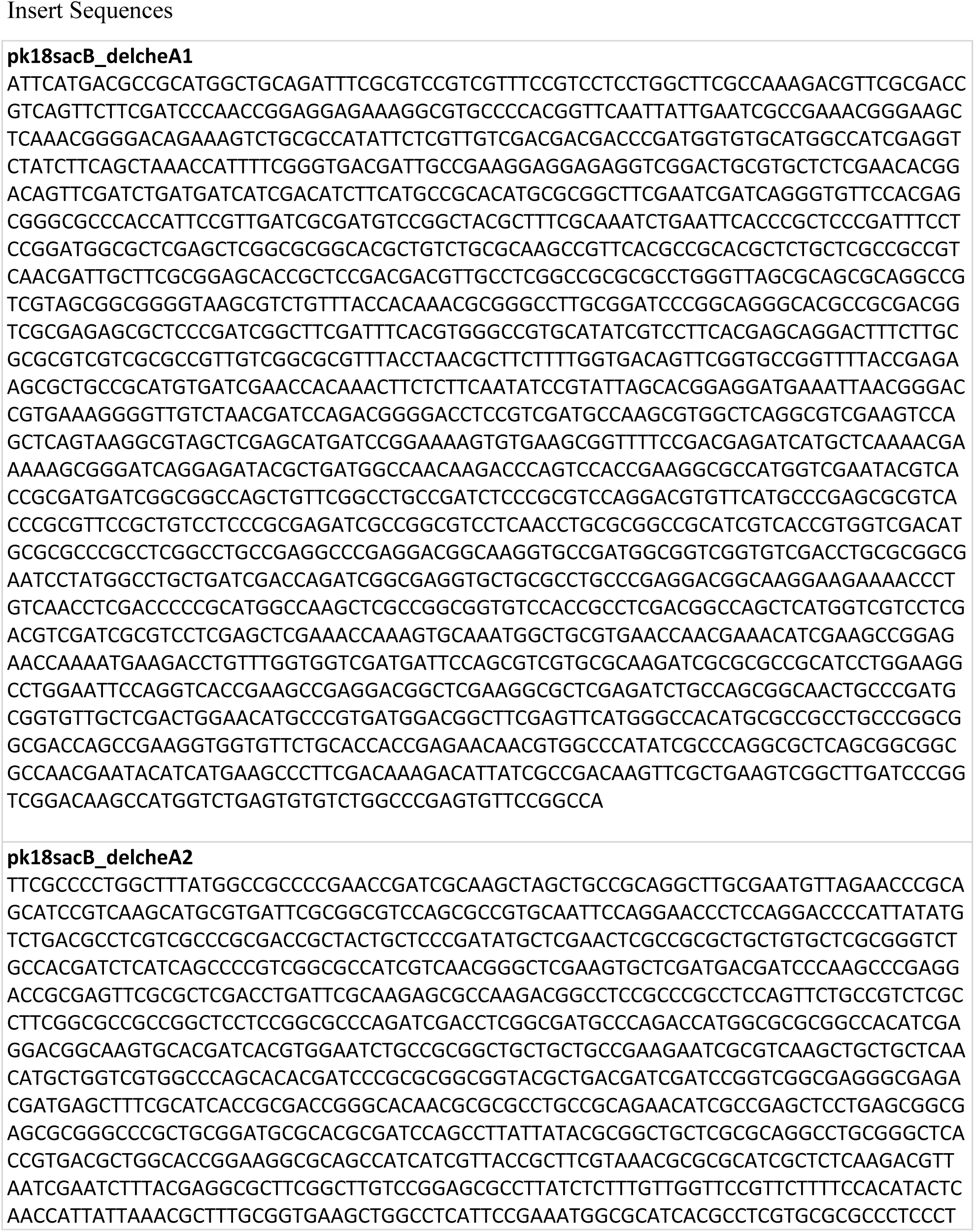
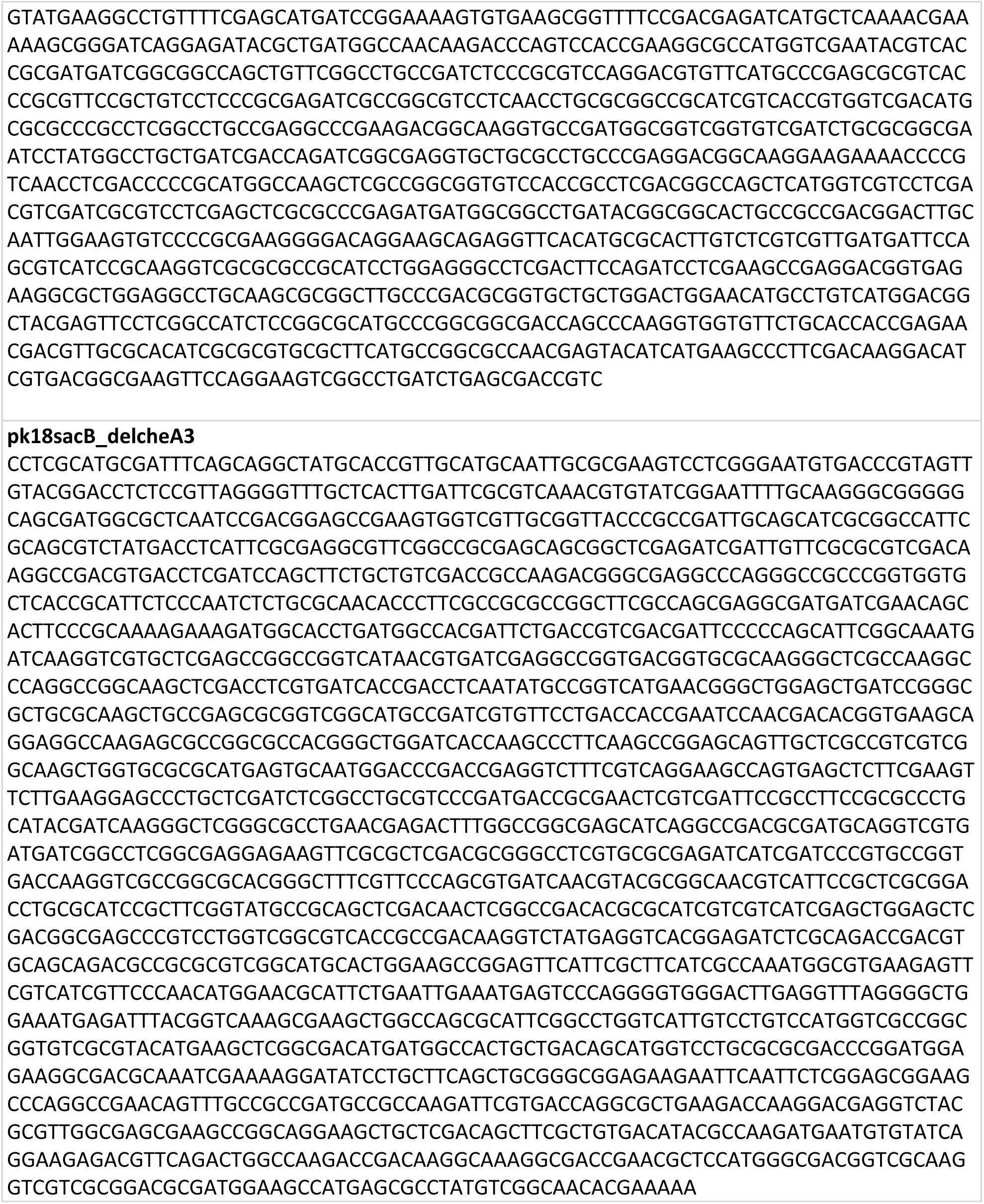
**Vectors Used for Homologous Recombination**. Insterted into XbaI cut site (vectors generated by Twist Biosciences)

**Table S1.**
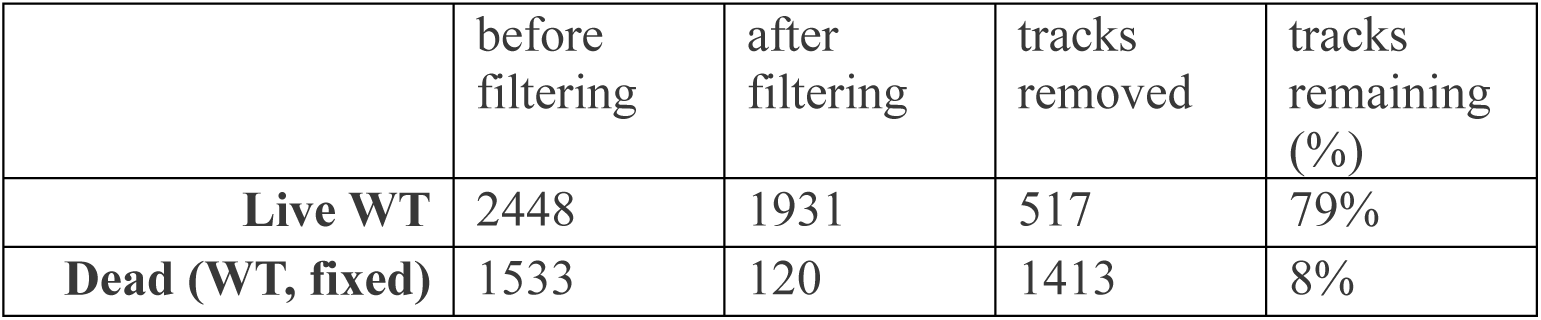
Filtering cutoff removes most dead cells but not most live cells.

**Table S1a.**
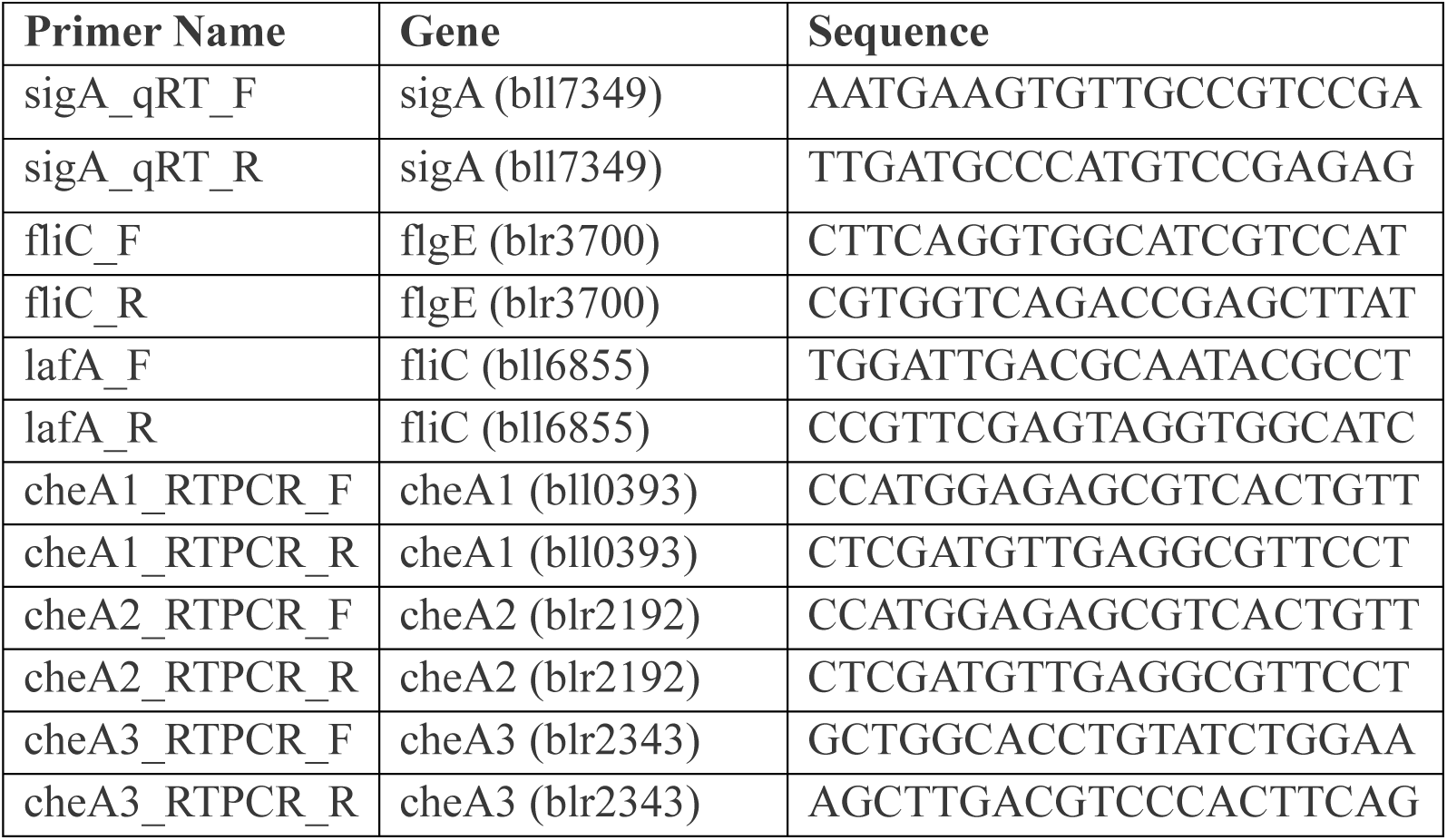
Primers used for qRT-PCR.

